# Subtractive Interaction Proteomics Reveal a Network of Signaling Pathways Activated by an Oncogenic Transcription Factor in Acute Myeloid Leukemia

**DOI:** 10.1101/464958

**Authors:** Nathalie Guillen, Maria Wieske, Andreas Otto, Afsar Ali Mian, Michal Rokicki, Carol Guy, Caroline Alvares, Paul Hole, Hannelore Held, Oliver Gerhard Ottmann, Dörte Becher, Marieangela Wilson, Kate J. Heesom, Martin Ruthardt, Claudia Chiriches

## Abstract

Acute myeloid leukemias (AML) are characterized by recurrent genomic alterations, often in transcriptional regulators, which form the basis on which current prognostication and therapeutic intervention is overlaid. In AML transformation can often be attributed to single chromosomal aberrations encoding oncogenes, such as t(15;17)-PML/RARα or t(6;9)-DEK/CAN but it is unclear how these aberrant transcription factors drive leukemic signaling and influence cellular responses to targeted therapies. Here we show that by using a novel “subtractive interaction proteomics” approach, the high risk AML-inducing oncogene t(6;9)-DEK/CAN directly activates signaling pathways that are driven by the ABL1, AKT/mTOR, and SRC family kinases. The interplay of these signaling pathways creates a network with nodes that are credible candidates for combinatorial therapeutic interventions. These results reveal specific interdependencies between nuclear oncogenes and cancer signaling pathways thus providing a foundation for the design of therapeutic strategies to better address the complexity of cancer signaling.

**Graphical Abstract:** 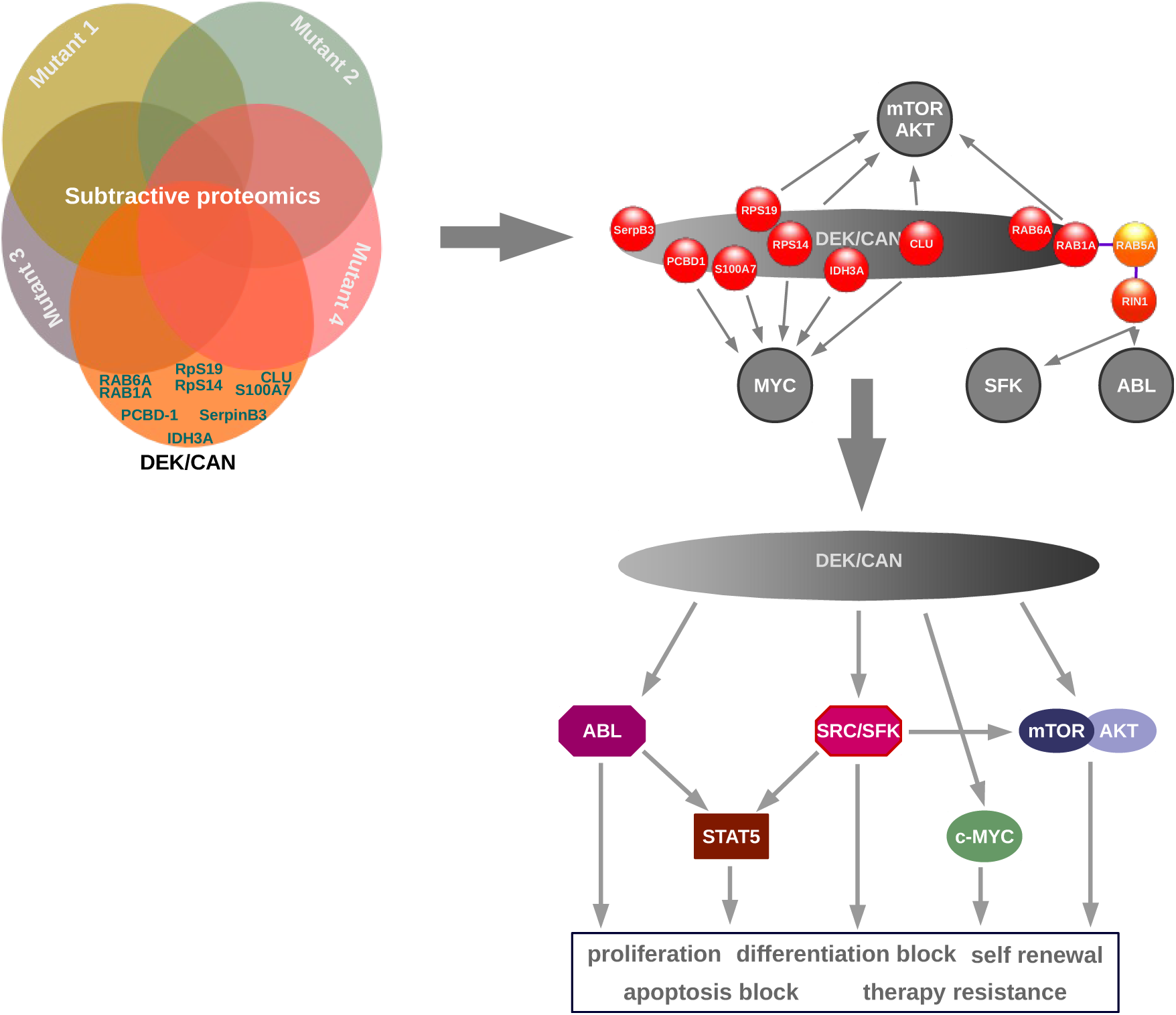

## Introduction

Cancer develops through the accumulation of somatic genomic alterations that act as driver mutations and determine the biology and the clinical behavior of the disease. Classification of cancers according to causal genetic mutations is exemplified by acute leukemias. In acute myeloid leukemia (AML), the WHO classification is based mainly on specific genomic aberrations and a detailed subclassification reflecting that the accumulation of mutational data derived from high-throughput sequencing approaches has linked genetic data with clinical management of AML. This led to a risk stratification based on mutational profiles for allocating patients to the best therapy option for definitive cure (Döhner et al., 2017; Gerstung et al., 2017; Grimwade et al., 2010; Papaemmanuil et al., 2016; Patel et al., 2012).

Analyses of the genomic landscape of AML has not only unraveled the complexity of the disease but also indicates pathogenetic mechanisms (Shlush et al., 2014; Welch et al., 2012), but translating the knowledge of the genetic and biologic intricacies of AML into clinical practice has been challenging. AML with specific, disease and risk defining karyotypes has allowed a better understanding of the molecular mechanisms of disease and recognition of the central role of aberrant transcription factors (class II mutations) in leukemogenesis. However, direct therapeutic targeting of transcription factor aberrations has been elusive and the most notable clinical success of treating t(15;17)-positive acute promyelocytic leukemia with ATRA and arsenic trioxide has not been initially based on rationale and molecularly driven design of therapy (Lo-Coco et al., 2013).

To date, most targeted therapies have focused largely on aberrant kinase signaling, caused by mutations e.g. of c-KIT or FLT3 and related signaling pathways, by combining kinase inhibitors with conventional chemotherapy. This approach has improved outcome, but has not evolved to curative therapy for AML; Stone et al., 2017). This may be due to the fact that these TKIs target signaling pathways that are linked to passenger but not to driver mutations; Papaemmanuil et al., 2016) and thus neither eradicate leukemic stem cells (LSC) nor overcome resistance to cytotoxic agents.

Three subtypes of AML carrying specific translocations, namely t(15;17), t(11;17) and t(6;9), are notable for being associated with a smaller number of co-existing driver mutations than for example AML with normal karyotype (Papaemmanuil et al., 2016; and our unpublished data). This strongly indicate that the function of the aberrant gene products may subsume the function of other driver mutations and thus does not necessarily require sequential acquisition of secondary genomic alterations.

Thus AML with the t(6;9) represent a perfect model for high risk AML. It is defined as a distinct entity by the WHO classification, because of its particular biological and clinical features and unmet clinical needs; Arber et al., 2016). In contrast to other AMLs (median age 66 years), most t(6;9)-AML patients are young, with a median age of 23-40 years (Appelbaum et al., 2006; Chi et al., 2008). Complete remission rates do not exceed 50% and median survival after diagnosis is only about 1 year (Grimwade et al., 2010; Tarlock et al., 2014). The t(6;9) represents the only structural chromosomal aberration present at diagnosis and encodes the DEK/CAN fusion protein; Oancea et al., 2014). Noticeably, the reciprocal CAN/DEK fusion transcript is not detectable (von Lindern et al., 1992), making DEK/CAN the unique gene product of t(6;9) providing convincing evidence for the decisive role of DEK/CAN in leukemia initiation.

The t(6;9)-DEK/CAN represents a class II mutation, which is leukemogenic; Oancea et al., 2010). Forced expression of DEK/CAN activates the AKT/mTOR pathway (Sandén et al., 2013) and STAT5 activation is part of the mechanism by which DEK/CAN induces leukemia; Oancea et al., 2014) suggesting that activation of several signaling pathways contribute to the specific features of t(6;9)-AML. However, the mechanisms by which DEK/CAN transforms cells and mediates therapy-resistance have not been completely resolved. While 30-75% of t(6;9)-AML patients harbor FLT3-ITD as a prognostically adverse risk feature at diagnosis (Oyarzo et al., 2004), FLT3-ITD accelerates the kinetics of relapse, but is not causally involved in leukemogenesis of t(6;9)-AML (Ommen et al., 2015 and our unpublished data). This is consistent with our findings that DEK/CAN transforms immature hematopoietic stem and progenitor cells (HSPCs) in which the FLT3 promoter is not active; Oancea et al., 2010). Accordingly, t(6;9) remains an independent adverse risk feature, irrespective of the presence of FLT3-ITD (Sandahl et al., 2014; Tarlock et al., 2014).

We therefore postulated the existence of a leukemogenic network in which the cooperating “synthetically” leukemogenic components did not require the presence of complementary mutations. By dissecting these multi-pathway mechanisms, we aimed to identify potentially fundamental mechanisms of signal pathway activation by class II mutations, and to shed light on the mechanisms of therapy resistance.

As this hypothesis was not amenable to purely genomic analysis, we reasoned that a DEK/CAN-mediated autonomous pathway activation should be reflected by its interactome. To analyze such a complex interaction network, we sought to achieve a maximal reduction of experimental variability by applying the novel strategy of “subtractive interaction proteomics” (SIP). This approach is based on a comparison of the interactome of an oncogene, with those of its functionally inactive mutants in the same genetic background in order to obtain eventually only relevant interaction partners. This is achieved by the subtraction of interaction partners that are common to its functionally inactive mutants classifying them as not relevant. We specified that the mutants to be used in SIP had to be biologically inactive but maintain the identical subcellular localization as the native factor to exclude modified interaction patterns due to an aberrant localization.

Here we present findings that are based on an in depth analysis of interactomes and related networks, which provides insights exceeding those provided by purely genetic informations. This will modify current concepts of pathogenesis of AML, mainly based on the two hit model postulating cooperation between class I and class II mutations which are considered independent in their contribution to leukemia induction (Deguchi and Gilliland, 2002).

## Results

### Putative DEK GSK3-phosphorylation sites and helices 3 and 5 of the CAN coiled coil domain (CC) are indispensable for the leukemogenic potential of DEK/CAN

Our preliminary structure-function analyses pointed to both CC domain in the CAN portion and putative phosphorylation sites in the DEK-portion, as indispensable structures for the functionality of DEK/CAN. Thus we mutated each structure in order to obtain the mutants necessary to run a complete SIP analysis with differential interactomes of DEK/CAN and these mutants.

In order to obtain functionally inactive CC-mutants we deleted each of 5 helices revealed in a 3D model of the putative CC domain in the CAN-portion created in I-TASSER (Figure 1A and S1A).

**Figure 1.**
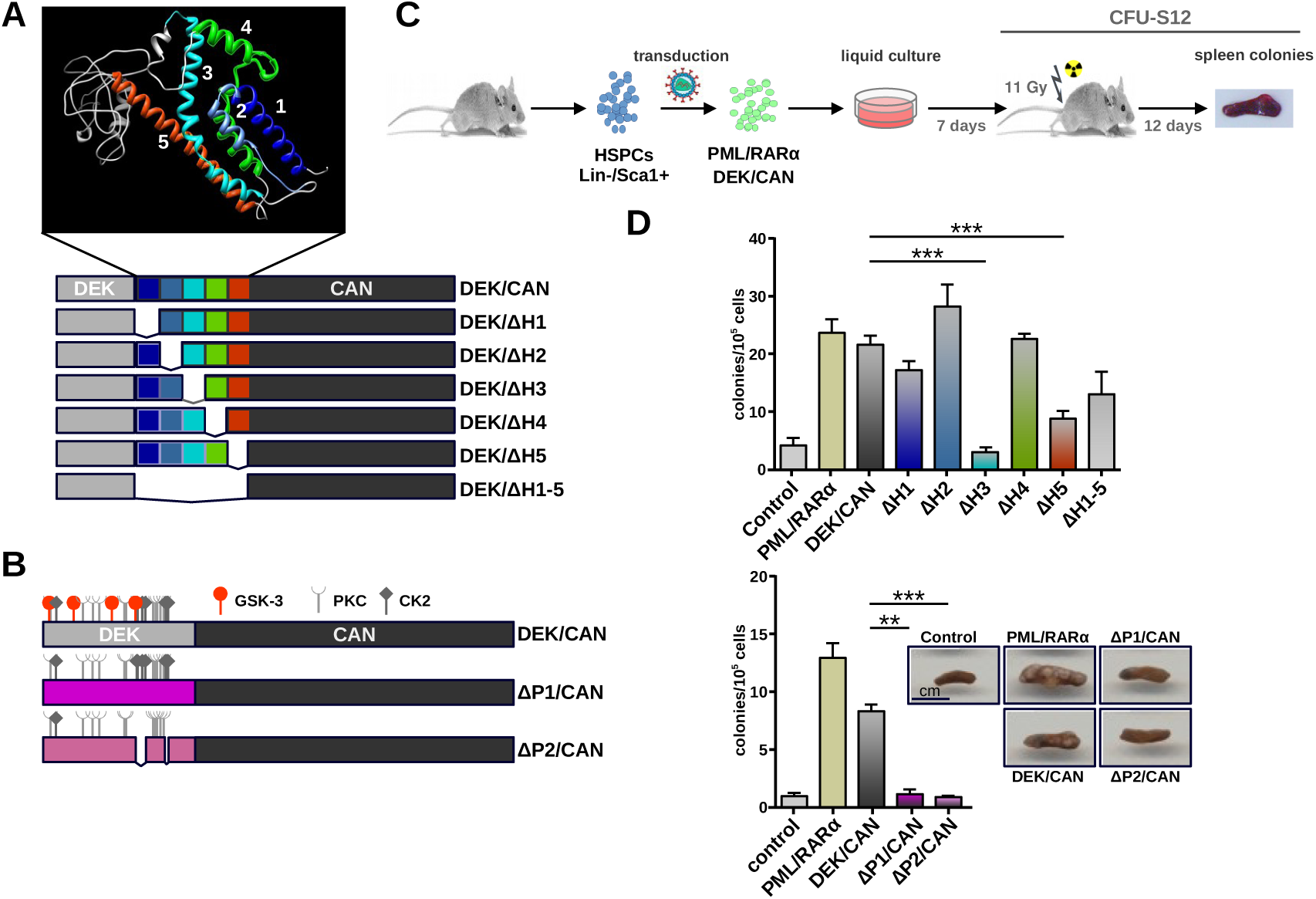
The leukemogenic potential of DEK/CAN mutants. **A.** I-Tasser 3D analysis of the DEK/CAN CC revealing 5 helices and modular organization of DEK/CAN’s helix-mutants (not in scale). **B.** Putative GSK3-, PKC-, and CKII-phosphorylation sites and modular organization of DEK/CAN’s phosphorylation mutants (not in scale). **C.** Influence of the mutations on the leukemogenic potential of DEK/CAN in murine Sca1+/lin-HSPCs. Cells were transduced and maintained for 7 days in liquid culture. 1×10^4^ cells were inoculated into lethally irradiated recipients. At day 12 spleen colonies were counted. **D.** Number of colonies in the spleens (n= 3). One representative experiment of three performed that yielded similar results is given (+/− SEM).

For the disruption of the functionality of the N-terminal DEK portion of the fusion protein we targeted first the putative GSK3-phosphorylation sites by point-mutations that changed Ser to Ala (ΔP1/CAN). In addition we also deleted a stretch of CK2 phosphorylation sites (ΔP2/CAN)(Figure 1B); Kappes et al., 2004). As a consequence overall phosphorylation of both mutant constructs was reduced compared to the native fusion protein (Figure S1B).

The loss of leukemogenic potential of these DEK/CAN mutants was investigated in a CFU-S12 assay (Figure1C); Oancea et al., 2010) on murine Sca1^+^/lin^−^ HSPC, that retrovirally expressed either DEK/CAN, its mutants or controls (empty vector, DEK/CAN, and PML/RARα). The deletion of helices 3 and 5 as well as the loss of the GSK3-phosphorylation sites alone and in combination with the deletion of CKII phosphorylation sites largely abrogated the functionality of DEK/CAN as revealed by the abolished colony formation in the spleen (Figure 1D).

In order to exclude that the loss of function was due to a disruption of normal sub-cellular localization we confirmed the co-localization of the four DEK/CAN mutants with DEK/CAN in 293T cells that co-expressed GFP-DEK/CAN either with HA-tagged ΔP1 and ΔP2/CAN or with SNAP-tagged DEK/ΔH3 and 5 (Figure S1C and D). These data confirmed that the four mutants fulfilled all criteria for subsequent SIP analysis.

### The subtractive interaction proteomics identifies nine exclusive DEK/CAN binding proteins

SIP requires an optimum enrichment of specific and relevant interaction partners. Tandem-affinity purification (TAP) is a two step protocol which allows a nearly complete purification of stable protein complexes under close to physiological conditions for subsequent analysis by tandem mass spectrometry (MS) (Bürckstümmer et al., 2006). Our TAP-tag encoded two Protein A sequences separated from a streptavidin-binding peptide (SBP) by a TEV cleavage site (Figure 2A). Any negative effect of this TAP-tag on the biological functionality of DEK/CAN construct was excluded in a CFU-S12 assays (Figure 2B). In order to further improve specificity, TAP-purification was combined with a SILAC strategy; Ong et al., 2002).

**Figure 2.**
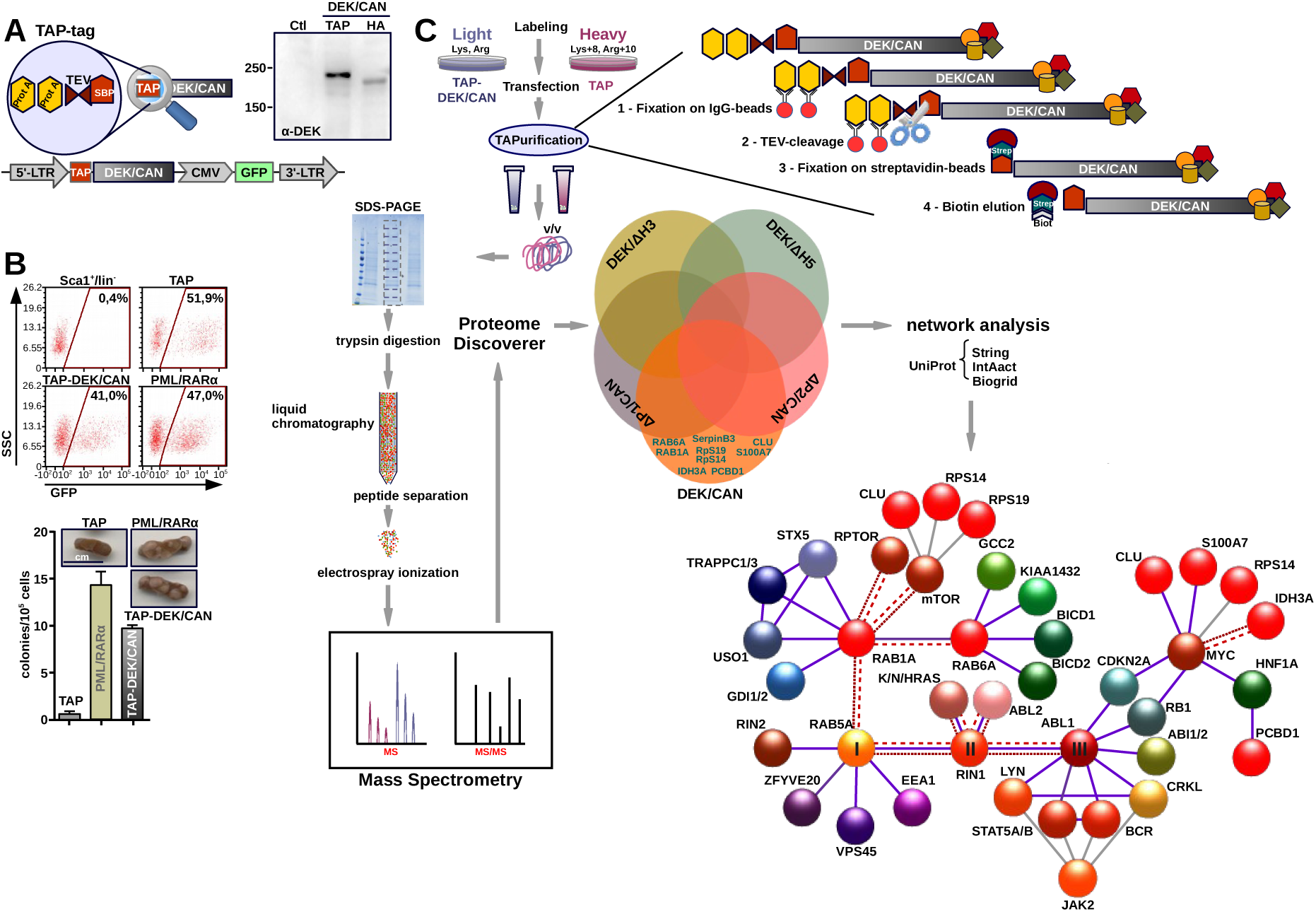
TAP-tag proteomics and SIP. **A.** Schematic representation of the TAP-tag construct. TAP - Tandem Affinity Purification; TEV – Tobacco Etch Virus protease; Prot A – protein A; SBP – Streptavidin Binding Protein. The provirus with TAP-tagged DEK/CAN used for the proteomics experiments (not in scale). The immunoblot shows the expression of TAP-tagged DEK/CAN in comparison to HA-tagged DEK/CAN in 293T cells (α-DEK staining). **B.** Leukemogenic potential of HSPCs transduced with the TAP-tagged DEK/CAN; PML/RARα - control. Transduction efficiency assessed by the detection of GFP. TAP – empty vector. **C.** Work flow of TAP-tag proteomics and tandem MS followed by data elaboration (Proteom Discoverer) and network analysis (BioGrid, IntAct, STRING). Synopsis of potential functional interaction network and signaling pathway involvement of DEK/CAN’s EBs. The Venn diagram shows only EBs. Functional network of the exclusive interactome of DEK/CAN - red spheres: DEK/CAN’s EBs. Evidence of potential functional interactions was obtained by STRING (purple lanes), Biogrid (dashed lines), and/or IntAct (dotted lines) or by data mining (grey lanes). The factors indicated with I, II, and III were taken as central for further analysis of potential functional interaction by STRING, Biogrid and/or IntAct analysis (see also Figure S2B-H and S3).

Therefore we transfected 293T cells that had previoulsy incorporated ^15^N_4_ ^13^C_6_-Arg and ^15^N_2_ ^13^C_5_-Lys (SILAC heavy - Svy) with the empty vector control (TAP), whereas those cultured in unmodified medium (SILAC light – Sli) were transfected with either the TAP-DEK/CAN or mutant constructs. After TAP-purification, the resulting protein complexes were appropriately prepared for an analysis by tandem MS. Peptide identification was perforemd by using Proteom Discoverer software (Figure 2C). Finally we made a list based on frequency of appearance and amount of positive labeled peptides (Table S1). Svy peptides corresponding to the TAP-control were subtracted as background noise.

Common binders (CBs) were Sli proteins which bound to both DEK/CAN and to at least one of the four mutants. By this we revealed already known interaction partners of either DEK, CAN or DEK/CAN, such as hCRM1 and verified the validity of our experimental approach (Table S1). By definition exclusive binders (EBs) interacted only with DEK/CAN after the subtraction of CBs from the total of DEK/CAN binders based on 2-7 replicates (Figure 2C; Table S1). Nine proteins fulfilled these criteria, namely Clusterin (CLU), RPS14 and 19, IDH3A, RAB1A, RAB6A, PCBD-1, S100A7 and SerpinB3 (Figure 2C). We confirmed the expression of these EBs in data sets of t(6;9) patients (n=3) from the UK-NCRI AML study group within the MILE study (Haferlach et al., 2010 and data not shown).

### The functional network analysis of DEK/CAN’s EBs predicts mTOR, ABL1, SFK and MYC as activated signaling pathways

We hypothesized that the set of EBs would be associated with signaling pathways related to t(6;9) induced leukemogenesis. Thus we interrogated three functional interaction databases integrated in the UniProt universal protein knowledge base (www.uniprot.org), STRING, BioGrid, and IntAct (Jensen et al., 2009; Kerrien et al., 2007; Stark et al., 2006) and dissected a network of signaling pathways related to these EBs by applying stringent parameters and the presence of EBs in at least two of three interrogated databases.

Interestingly a close interaction between RAB1A and RAB6A, both EBs, as well as between RAB1A and RAB5A, which was not an EBs but a CB (Table S1), was revealed. Also interaction with mTOR and RAPTOR was shown in accordance to RAB1A’s known role in the activation of this pathway; Thomas et al., 2014) (Figure 3A). A further refinement of the subnetworks of RAB1A/RAB6A/RAB5A by STRING and Biogrid analyses identified a functional interaction with ABL1 kinase (Figure 2C), not known yet, to be associated with t(6;9)-AML; Balaji et al., 2012). Further analysis revealed functional interactions with STAT5, STAT3 and JAK2, all known to be activated in syngeneic DEK/CAN-AML cells; Oancea et al., 2014). Notably, the SRC family kinase (SFK) LYN appeared a highly confident functional interaction partner embedded in the signaling network (Figure 2C). In addition, a direct functional relationship between the EBs CLU, S100A7 and c-MYC, in the case of PCBD-1 mediated by HNF1A, was seen (Figure 2C). These analyses together with in depth data mining indicated the involvement of DEK/CAN’s EBs and thereby DEK/CAN itself in the activation of the following signaling pathways (Figure 3A): AKT/mTOR (Jo et al., 2008; Payne et al., 2012; Thomas et al., 2014; Yip et al., 2013), SFK, ABL1 (Cao et al., 2008; Ting et al., 2015), and c-MYC (Deol et al., 2011; Fröjmark et al., 2010), all known to be involved in the pathogenesis of cancer, maintenance of cancer stem cell, and determination of therapy response and therefore referred to as cancer activated signaling pathways (CASPs).

**Figure 3.**
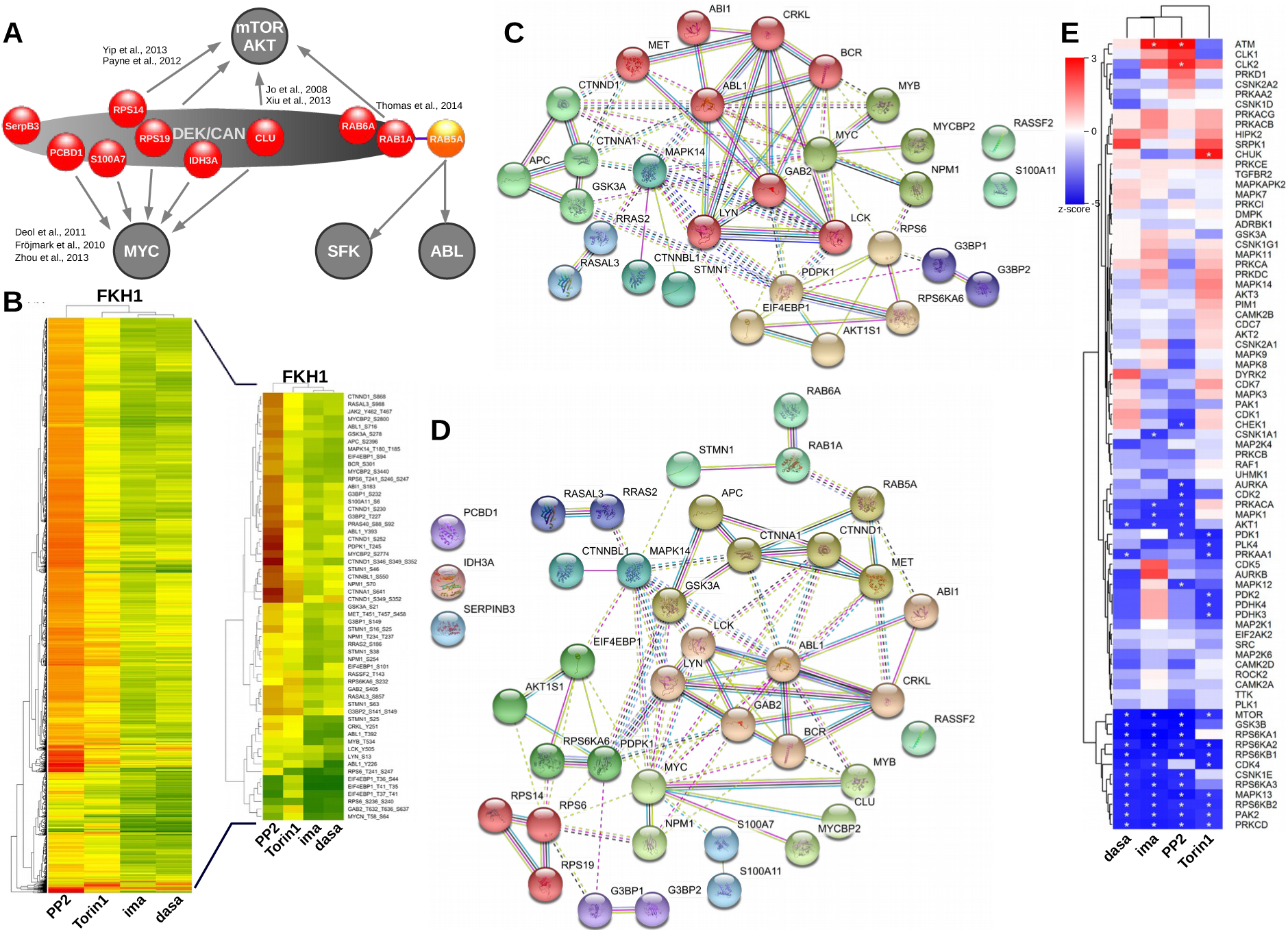
Predicted signaling pathways regulation by DEK/CAN’s EBs. **A.** Relationship between EBs and potentially activated signaling pathways. **B.** Comparative phosphoproteomics on FKH1 treated to target predicted signaling pathways: PP2 - SFK, Torin1 - AKT/mTOR, imatinib - ABL1, dasatinib - ABL1/SFK. The heatmap represents the fold up-(red) or downregulation (green) of single phosphopeptides of treated vs untreated cells. Fold changes of phosphopeptides representing key factors in the predicted signaling pathways were extracted. **C-D.** STRING cluster analysis of key factors in the predicted signaling pathways **C.** first alone **D.** and then in combination with DEK/CAN’s EBs. Line colors indicate the kind of evidence the functional interaction is based on – magenta – experimental; cyan – from curated databases; yellow – data mining; black – co-expression; red - gene fusion; blue – gene co-occurence; purpur – protein homology. **E.** Effect of treatment on master kinases – KSEA. The comparative phosphoproteomics were analyzed using KSEA based on PSP and NetworKIN (score > 5) with a cut-off of at least 3 substrates for the identification of a master kinase; Wiredja et al., 2017).

These CASPs are regulated by phosphorylation cascades. Therefore as a first step of validation we performed a phosphoproteomic screen on the t(6;9)-positive patient derived cell line FKH1. Based on predicted signaling pathway profile we exposed these cells to inhibitors of SFK (PP2), mTOR/AKT (Torin1), ABL1 (imatinib) and ABL1/SFK (dasatinib). Fold changes in single phosphorylated sites are given as heatmap of about 3.500 phosphopeptides based on the ratio between untreated (set at 1) and treated cells (Figure 3B). The way how inhibitors influence these phosphopeptides and related signaling pathways was analyzed by Ingenuity Pathway Analysis (IPA) software (Quiagen). Noteworthy both Torin1 and PP2 seemed to work by an increased phosphorylation of regulators whereas imatinib and dasatinib led to a reduction of overall phosphorylation of targets (Figure S4). A cluster and network analysis of factors known to be involved in ABL1-, mTOR/AKT-, SFK-, and MYC signaling by STRING revealed a significant functional interaction between all about 30 factors with the exception of two (Figure 3C). In addition, 4 major clusters (>3 components) were revealed, which were focused on GSK3, ABL1 and SFK (LYN and LCK), mTOR/AKT (AKT-S1 component of the mTOR complex) and MYC. The introduction of the EBs into this cluster analysis further confirmed their close functional relationship with these CASPs (Figure 3D).

Kinase substrate enrichment analysis (KSEA) allows to systematically infer the activation of kinases based on a profile of phosphorylated substrates obtained by MS-based phosphoproteomics; Casado et al., 2013). We used this approach to understand which kinases were affected by the exposure of FKH1 cells to the above mentioned inhibitors. KSEA uses the PhophoSitePlus (PSP) and the NetworKin databases. PSP annotations are restricted to human proteins and the NetworKin confidence score (score > 5) allows to adjust the inclusion stringency with a cut-off of at least 3 substrates for the identification of a master kinase; Wiredja et al., 2017). Interestingly, all inhibitors inhibited significantly the mTOR cascade suggesting mTOR to be a master kinase and its signaling cascade central for the DEK/CAN induced leukemic phenotype (Figure 3E).

### t(6;9)-DEK/CAN determines AKT/mTOR signaling activation

The SIP based prediction of a relationship between DEK/CAN and AKT/mTOR confirmed by KSEA revealing mTOR a master kinase in FKH1, prompted us to investigate the role of AKT/mTOR signaling for t(6;9)-AML. First, we studied AKT/mTOR signaling in FKH1 cells by ICF for pAKT-Ser473 which revealed a strong activation of AKT in these cells (Figure 4A).

**Figure 4.**
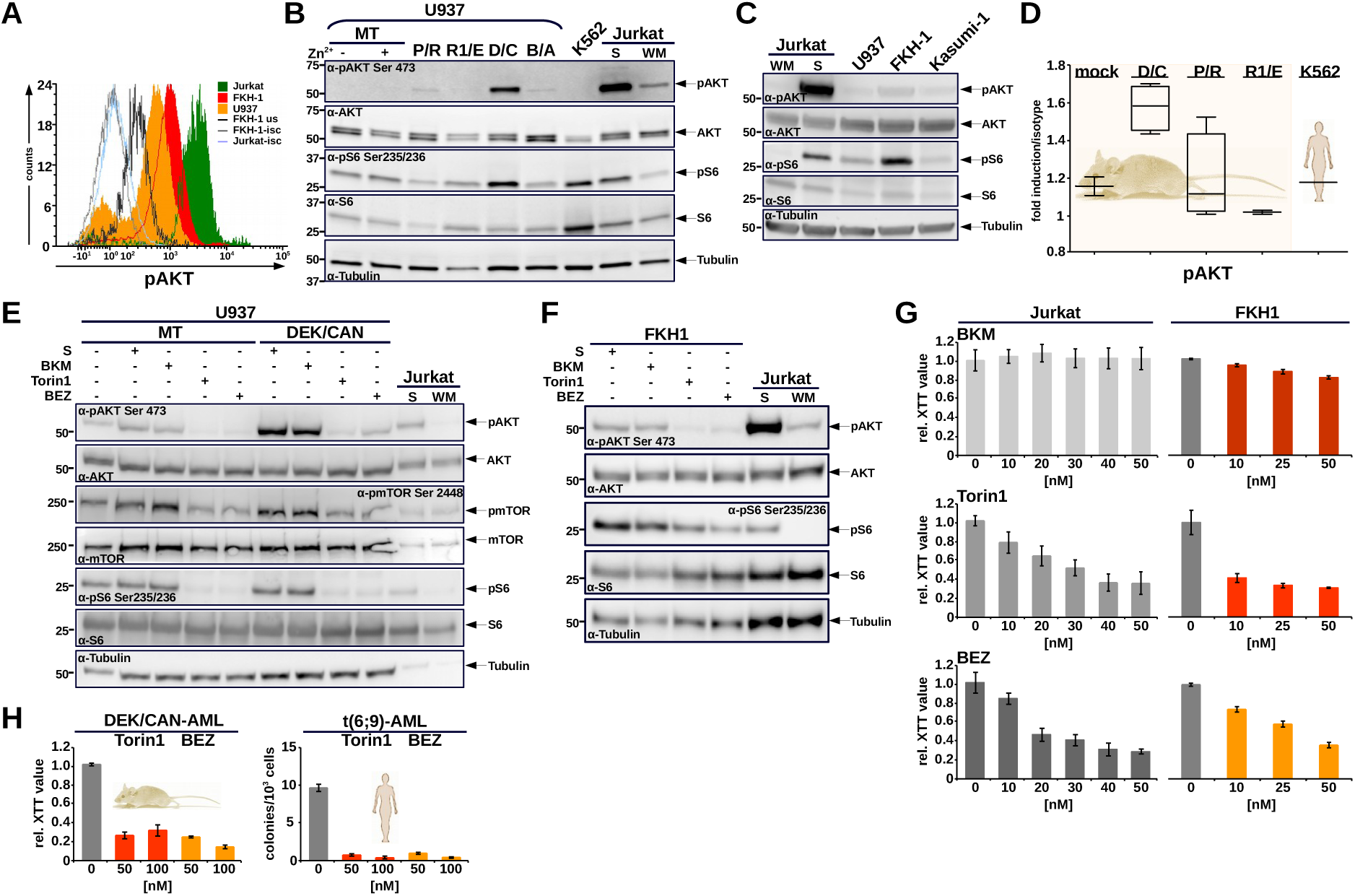
mTOR/AKT signaling in t(6;9)-AML. **A.** Detection of phosphorylated AKT (Ser473) in FKH1 by ICF. U937, Jurkat – negative and positive controls. **B.** Activation of mTOR/AKT in U937 expressing transgenes P/R – PML/RARα; R1/E – RUNX1/ETO; D/C – DEK/CAN; B/A – BCR/ABL under the control of a Zn^2+^-inducible metallothionein (MT) promoter. U937 MT +/− Zn^2+^ - empty vector; K562, Jurkat - positive controls. S – solvent; WM – wortmannin. The blots were stained with the indicated antibodies. Tubulin – loading control. **C.** mTOR/AKT signaling in patient derived AML cells. U937, Jurkat – negative and positive controls, S – solvent; WM – wortmannin; FKH1 – t(6;9)-DEK/CAN; Kasumi1 – t(8;21)-RUNX1/ETO. **D.** mTOR/AKT signaling in syngeneic AMLs. Mock – BM cells from healthy mice transplantated with empty vector transduced HSPCs; D/C, P/R, and R1/E – blast from syngeneic leukemias driven by DEK/CAN, PML/RARα, or RUNX1/ETO, respectively. K562 – positive control. **E.** PI3K/AKT/mTOR inhibition in U937 expressing DEK/CAN. All indicated inhibitors were used at 50nM. Jurkat – positive controls, S – solvent; WM – wortmannin. **F.** PI3K/AKT/mTOR inhibition in FKH1. Jurkat – positive control, S – solvent; WM – wortmannin. All inhibitors were used at 50nM. **G.** PI3K/AKT/mTOR inhibition in FKH1 - proliferation. FKH1 and Jurkat (control) were exposed to the indicated dosages of inhibitors. Proliferation/cytotoxicity was measured by XTT. The data represent the mean of two experiments +/− SEM. **H.** PI3K/AKT/mTOR inhibition in syngeneic DEK/CAN-positive and human t(6;9)-AML cells - proliferation (XTT) or CFU formation. The indicated dosages of Torin1 or BEZ were used. The data represent the mean of two experiments +/− SEM.

The comparison of the AKT/mTOR activation by t(15;17)-PML/RARα (P/R), t(8;21)-RUNX1/ETO (R1/E) and t(9;22)-BCR/ABL (B/A) with that by DEK/CAN (D/C) in U937 showed that this activation was specifically related to the expression of t(6;9)-DEK/CAN and not a common feature of AML specific translocations. Slight AKT activation was seen only in P/R-and B/A-but not in R1/E-harboring U937 cells. DEK/CAN (D/C) potently promoted AKT/mTOR axis as confirmed by a strong pS6 signal, one of the down-stream targets of AKT/mTOR signaling (Figure 4B). The applicability of these observations to patient derived leukemic cells inherently harboring the DEK/CAN or RUNX1/ETO, respectively, was confirmed in FKH1 and Kasumi-1 cells where a strong pAKT/pS6-phosphorylation was found in FKH1, but only a minimal pAKT signal in Kasumi-1 cells (Figure 4C). As third line of evidence for the DEK/CAN mediated AKT/mTOR activation we demonstrated that syngeneic DEK/CAN-AMLs exhibited a higher pAKT signal as compared to PML/RARα-or RUNX-1/ETO-induced AMLs (Figure 4D).

The mechanisms by which DEK/CAN activates the AKT/mTOR cascade has not been elucidated yet. Activation of AKT has been attributed to either receptor induced PI3K signaling or autonomously by mTOR complex 2 (mTORC2); Lopiccolo et al., 2007). To dissect the level at which the AKT/mTOR signaling is activated we used three pharmacological inhibitors with distinct inhibitory properties: Torin1, a combined mTORC1/C2 inhibitor, NVP-BEZ235 (BEZ), a dual PI3K and mTORC1/C2 inhibitor, and NVP-BKM120 (BKM), a selective PI3K inhibitor; Badura et al., 2013). Only those compounds with mTORC2 inhibiting activity (Torin1 and BEZ) suppressed the activation of the AKT/S6K cascade in FKH1 cells (Figure 4E and F). Interestingly, PI3K-inhibition did not have any effect on the activation of AKT itself.

AKT/mTOR activation is a central mediator of cell survival, cell growth and proliferation (Laplante and Sabatini, 2012). Here we exposed FKH1 cells, syngeneic DEK/CAN-AML and primary t(6;9)-AML blasts to BKM, BEZ, and Torin1 to determine if proliferation of DEK/CAN-positive leukemic cells is dependent on AKT/mTOR activation. While inhibition by Torin 1 and BEZ, efficiently arrested growth at low concentrations, the selective PI3K inhibitor BKM had no effect on leukemic cell growth (Figure 4G). Moreover proliferation of syngeneic DEK/CAN-AML cells and colony formation of t(6;9)-AML blasts were strongly reduced upon exposure to Torin1 or BEZ (Figure 4H).

These data show that the activation of the AKT/mTOR axis is indispensable for the growth of t(6;9)-DEK/CAN positive leukemic cells and independent of PI3K confirming its central role for t(6;9)-AML.

### ABL1-kinase is activated in DEK/CAN-driven leukemic cells

Our SIP data suggested an ABL1 activation in t(6;9)-AML by a functional interaction between DEK/CAN and ABL1 mediated by RAB1A/RAB5A/RIN1 (Figure S3G-J). Here we definitively confirmed ABL1 activation in FKH1 cells by ICF investigating both ABL1 autophosphorylation at Y412 and substrate-phosphorylation of CRKL. In addition we assessed the effect of ABL1-directed kinase inhibitors imatinib or dasatinib which led to dephosphorylation of ABL1 as well as of its substrates CRKL (Figure 5A-C) and STAT5 (Figure 5D). To exclude a FKH1 related cell line effect we compared autophosphorylation of ABL1 in DEK/CAN-with that in BCR/ABL1-driven syngeneic leukemia cells. Both BCR/ABL1-and DEK/CAN-positive leukemia cells exhibited the same phospho-ABL speckled cytoplasmic staining pattern as in human FKH1 and K562 cells (Figure 5E).

**Figure 5.**
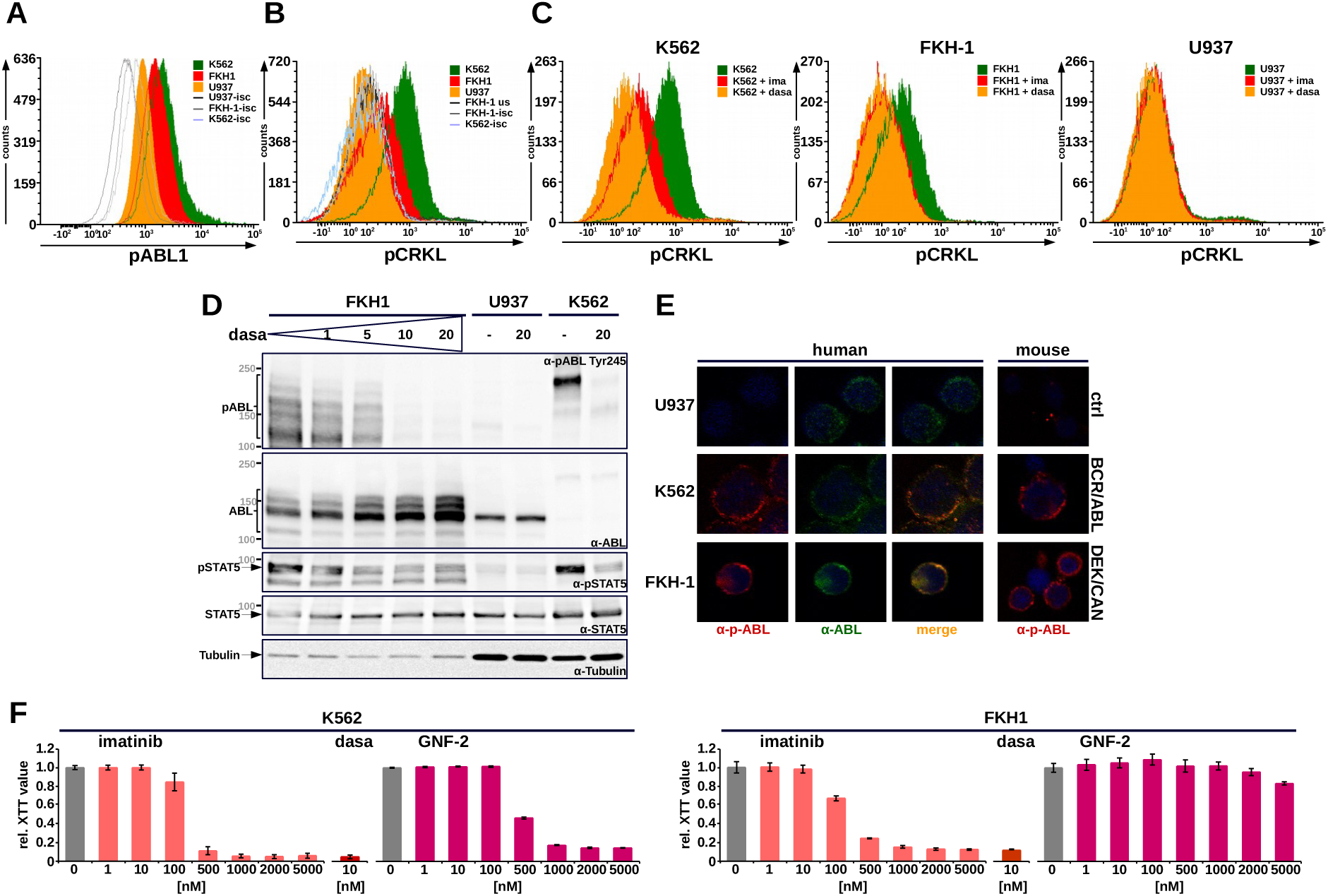
ABL1 activitation in t(6;9)-AML. **A.** pABL1 (Y412) in FKH1 cells - ICF. U937, K562 negative and positive control. **B-C.** ABL1 substrate activation in FKH1 – ICF of pCRKL; ima(tinib); dasa(tinib). **D.** ABL1-STAT5 signaling in FKH1 – immunoblotting. The blots were stained with the indicated antibodies. **E.** Immunofluorescence staining pattern of ABL1 in human and murine DEK/CAN-AML cells. K562, syngeneic CML like disease - positive controls. U937, murine HSPCs - ctrl. **F.** Allosteric inhibition of ABL1 by GNF-2 in comparison to imatinib in FKH1. Proliferation/cytotoxicity was measured by XTT. The bars represent means of three independent experiments each performed in triplicates +/− SD. K562 - positive controls.

ABL1 activation can be attributed to a variety of different mechanism such as SH2-, SH3-and/or myristoyl binding pocket (MBP)-relocalization (Hantschel et al., 2003). The latter can potentially be inhibited by allosteric MBP mimics, a class of drugs that are efficiently inhibiting activated ABL1 kinases and have now entered clinical trials (Wylie et al., 2017). To study the impact of MBP relocation as a mechanism of activation in DEK/CAN-positive leukemic cells, we exposed FKH1 and K562 cells to GNF-2, an efficient allosteric ABL1 inhibitor (Adrián et al., 2006). GNF2 fully inhibited K562 cells, but not FKH1 cells, in contrast to imatinib or dasatinib (Figure 5F). Thus the DEK/CAN mediated ABL1 activation is not subject to allosteric inhibition through the occupancy of the MBP which is consistent with previous data showing that RIN1 mediated activation occurs via the SH2-or SH3 displacement; Hu et al., 2005).

### SRC family kinases are active in t(6;9)-AML

The activation of SFK such as SRC or LYN predicted by our network analysis of DEK/CAN’s EBs was confirmed in FKH1 (Figure 6A). The significance of SFK activation for the t(6;9)-AML was shown by the fact that the exposition to dasatinib and ponatinib, both potent SFK inhibitors, resulted in dephosphorylation of SRC and LYN as well as STAT5 (Figure 6A and S5A). This was accompanied by a dose dependent reduction of proliferation in FKH1 and syngeneic DEK/CAN-AML cells (Figure 6B and C and S5B) as well as by a reduced CFU formation in both human t(6;9)-AML blasts and syngeneic DEK/CAN-AML cells (Figure 6D and E).

**Figure 6.**
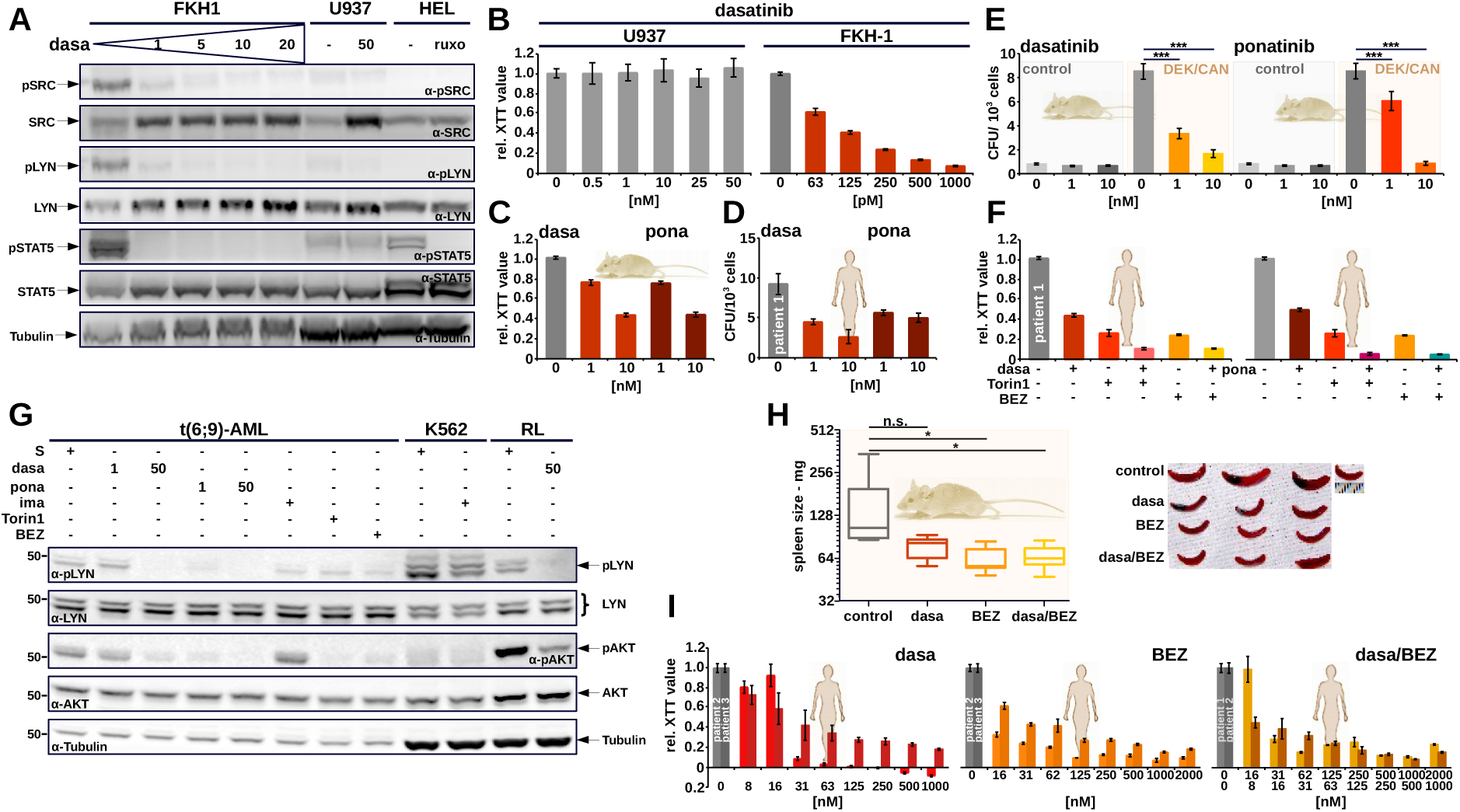
SRC family kinase activation in t(6;9) positive AML. **A.** SFK inhibition in FKH1. Dasa(tinib) - SFK inhibitor. The indicated antibodies detected phosphorylated and total SRC, LYN, and STAT5. Tubulin – loading control. U937, HEL and RL cells - controls. **B-D. SFK inhibition in t(6;9)-AML. B.** Proliferation in FKH1 - relative XTT values represent the mean of three independent experiments +/− SEM. **C.** Syngeneic DEK/CAN-leukemic cells exposed to dasa(tinib) or pona(tinib). Relative XTT values of one representative experiment of three each performed in triplicate +/− SEM yielding similar results are shown. **D.** SFK inhibition in human t(6;9)-AML. CFU assay in semi-solid medium. Data represent one representative experiment of two, each performed in triplicates +/− SEM. **E.** SFK inhibition in syngeneic DEK/CAN-AML – CFU-assay on syngeneic blasts. Control – BM cells from healthy mice transplantated with empty vector transduced HSPCs. Data represent the mean of three experiments each performed in triplicates +/− SD. *** p<0.001. **F-H. Crosstalk between SFK and AKT/mTOR signaling in t(6;9)-AML. F.** SFK/ABL1 and AKT/mTOR inhibition in human t(6;9)-AML cells. Relative XTT values from one representative experiment of two yielding similar results and each performed in triplicate +/− SEM. **G.** Immunoblotting stained with the indicated antibodies for SFK and AKT/mTOR activation. Dasa(tinib) and pona(tinib) were used at the indicated dosage, ima(tinib) −1μM, BEZ and Torin1 – 50nM. K562 and RL cells – controls. Tubulin – loading control. **H.** SFK/ABL1 and AKT/mTOR inhibition in syngeneic DEK/CAN-AML *in vivo* – spleen size. Control - solvent (n=7), dasa (n=7), BEZ (n=8), BEZ/dasa (n=8). Three spleens for each group in comparison to one spleen from a healthy mouse as reference are shown.

The fact that KSEA revealed AKT/mTOR signaling as regulated by SFK inhibitors prompted us to investigate the relationship between AKT/mTOR and SFK activity. We used dasatinib and ponatinib as SFK inhibitors, and Torin1 and BEZ as AKT/mTOR inhibitors on primary t(6;9)-AML blasts. Both SFK and AKT/mTOR inhibitors alone strongly reduced but the combination of the two classes abolished growth of t(6;9)-AML blasts nearly completely (Figure 6F). Interestingly, SFK inhibition led to reduced AKT activation, whereas inhibition of AKT/mTOR did not have any effect on SFK activation, suggesting that SFK are up stream regulators of AKT/mTOR activity. Imatinib, as selective ABL1 inhibitor, did not show any effect on either pathways (Figure 6G). These findings were confirmed by their effect on leukemogenic potential of DEK/CAN, as shown by the reduction of spleen size in DEK/CAN-AML mice (Figure 6H) and on blasts of two additional t(6;9)-AML patients upon treatment with dasatinib, BEZ or their combination (Figure 6I).

### In t(6;9)-AML STAT5 activation is regulated by a network of collaborating factors independently of JAK2

Profound activation of STAT5 is a known characteristic feature of t(6;9)-AML shown to be independent of FLT3-ITD; Oancea et al., 2014). This was related to the expression of DEK/CAN in both human FKH1 and in syngeneic DEK/CAN-AML cells (Figure 7A and B) and highly significant for t(6;9)-AML. In fact, AZD1208, a pharmacological inhibitor of PIM1/2 kinases, downstream effectors of activated STAT5; Peltola et al., 2004) induced a dose-dependent cell growth inhibition in FKH1 (Figure 7C). Although the network analysis clearly showed a functional link between ABL1, SFKs, and STAT5 in relationship to DEK/CAN’s EB, the enrichment of phosphopeptides for phosphoproteomics does not include phosphorylated STAT5, for still unknown reasons. As the lack of activity of ruxolitinib on FKH1 in contrast to JAK2-V617F-dependent HEL cells suggested, STAT5 activation on t(6;9) was independent of JAK2 (Figure S5C). On the other hand we show that STAT5 activation was regulated by an interplay between AKT/mTOR, SFK ad ABL1. Intriguingly, inhibition by BEZ led to an up-regulation of phosphorylated STAT5 in FKH1 cells, which was not influenced by ruxolitinib as confirmed by the lack of any combined effect of JAK2 and AKT/mTOR inhibition on cell growth (Figure 7D and E).

**Figure 7.**
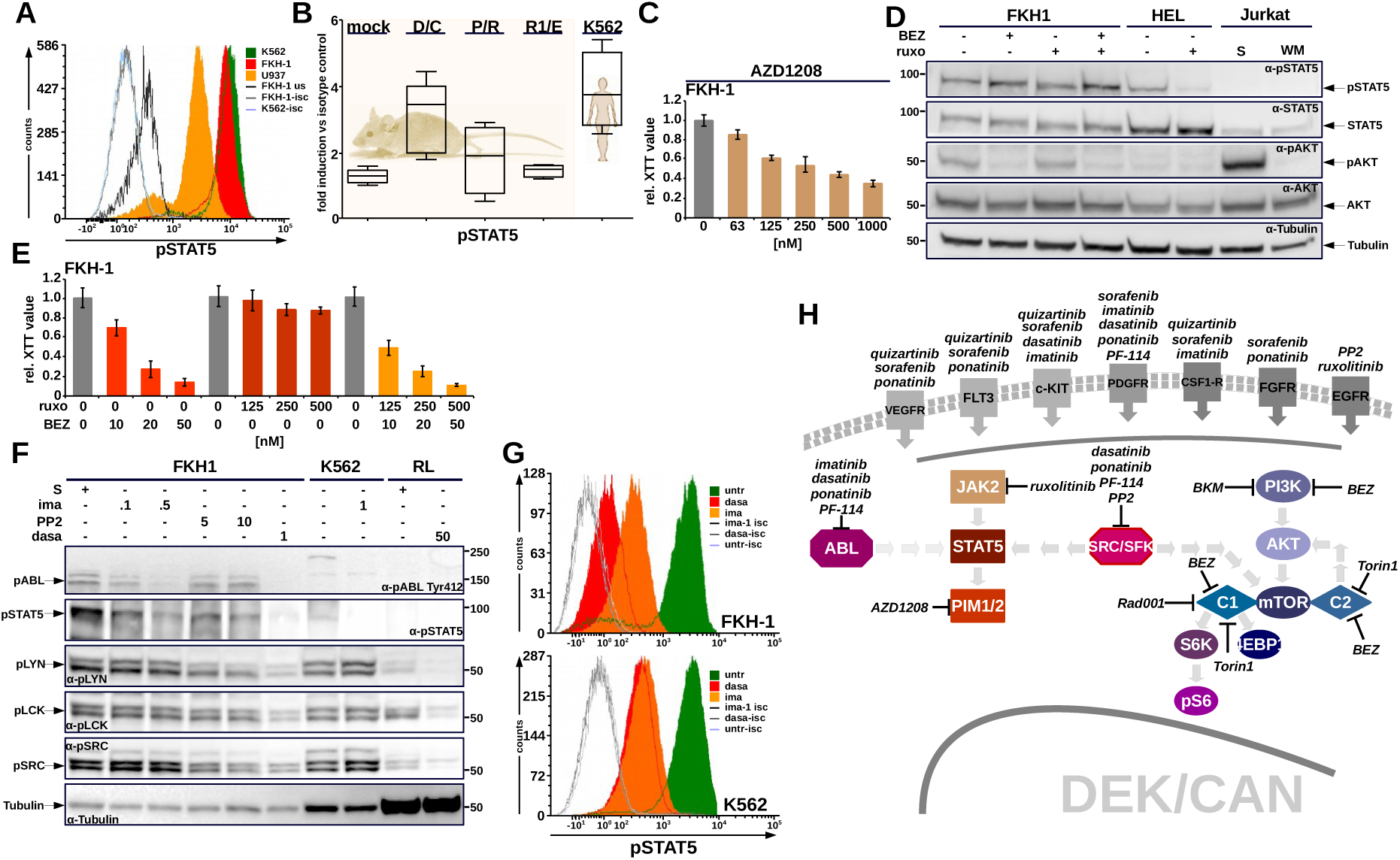
Regulation of STAT5 activation in t(6;9)-AML. **A.** STAT5 activation in FKH1 – ICF. U937, K562 - negative and positive controls. **B.** STAT5 activation in syngeneic AMLs. Mock – BM cells from healthy mice transplantated with empty vector transduced HSPCs; D/C, P/R, and R1/E – blast from syngeneic leukemias driven by DEK/CAN, PML/RARα, or RUNX1/ETO, respectively. K562 – positive control. **C.** PIM1/2 inhibition in FKH1 - AZD1208. Relative XTT values of three independent experiments +/−SD are given. **D.** Relationship between JAK2, mTOR/AKT and STAT5 – immunoblot. FKH1 were exposed to 50nm BEZ and 500nM ruxolitinib (ruxo) alone and in combination. The indicated antibodies were used for the detection of phosphorylated and total STAT5 and AKT. Tubulin – loading control. HEL, Jurkat - controls. **E.** JAK2 and mTOR/AKT inhibition in FKH1 cells alone and in combination. Relative XTT values represent the mean of three independent experiments each performed in triplicates +/− SD. **F.** Role of SFK and ABL1 for STAT5 regulation – immunoblot. FKH1 were exposed to the indicated concentrations (ima – μM) PP2 (nM) and 1 nM dasatinib. The indicated antibodies were used for the detection of phosphorylated ABL1 STAT5, and AKT. Tubulin – loading control. **G.** Contribution of SFK and ABL1 in STAT5 regulation – ICF. FKH1 and K562 (control) were treated with dasa (ABL1 and SFK) and ima (ABL1). **H.** Summary of the signaling network activated by the presence of DEK/CAN.

To understand the role of SFK and ABL1 for the regulation of STAT5 activation we used dasatinib, a dual ABL1/SFK inhibitor, and selective inhibitors of SFK (PP2) and ABL1 (imatinib) in order to address relative contribution of SFK and ABL1 to the STAT5 activation in t(6;9)-AML. As expected, imatinib resulted in dephosphorylation of ABL1 but not of SRC, LCK or LYN, while PP2 inhibited SFKs but not ABL1.

Conspicuously, both inhibitors resulted in partial inhibition of STAT5 whereas dasatinib completely dephosphorylated STAT5 (Figure 7F). ICF confirmed that STAT5 activation was dependent on both ABL1 and SFK activation in FKH1, whereas in K562 STAT5 activation could be solely attributed to the kinase activity of BCR/ABL (Figure 7G).

Taken together these data identify SFK and ABL1 as independent nodes which cooperate in regulation of STAT5, which involves also AKT/mTOR signaling.

## Discussion

Here we demonstrate the potential of interactome analysis of a leukemogenic transcription factor to uncover novel signaling networks involving its interaction partners. We provide clear evidence that DEK/CAN, a class II mutation, is directly involved in CASPs and such a critical relationship between kinase activation patterns and a specific genetic aberration represent a paradigm shift from AML seen as a transcription factor driven disease towards a kinase driven disease.

We show the autonomous activation of kinase activity by a class II mutation. This implied that inhibition of receptor tyrosine kinases (RTKs) would have no impact on their activation. In fact we excluded a contribution of the major RTKs known to be involved in activation of AKT/mTOR, SFK, and ABL1 (Figure S6A and B). Thus we propose a mechanistic model based on the aberrant localization of DEK/CAN to nuclear speckles mediated by the DEK-portion of the fusion protein. The consequences of this delocalization affect both CAN-binding proteins, transposed to a new environment, as well as the DEK-interactome, which is in juxtaposition with novel protein complexes which it does not normally encounter. This leads to functional changes of interaction partners, which conceivably would change into loss or gain of function. For other class II mutations such as PML/RARα and RUNX1/ETO such a sequester has been shown e.g. for HDAC; Grignani et al., 1998) or VDR; Puccetti et al., 2002).

Sequestration and thus inactivation of RPS14 and 19 by DEK/CAN supports a functional link between DEK/CAN and AKT/mTOR activation. The knockdown of RPS14 and RPS19 in zebrafish and human CD34+ cells leads to AKT/mTOR activation; Payne et al., 2012). Furthermore RPS14 and RPS19 haploinsufficiency has an important pathophysiological role in MDS and Diamond-Blackfan anemia (DBA), respectively; Ebert et al., 2008), where notably, mTOR activation is a common feature.

Our novel finding of ABL1-and SFK-activation by DEK/CAN was predicted by a complex formation between RAB1A/RAB6A via RAB5A involving RIN1, a known activator of ABL1. Additional multi-layered network analysis revealed a link of this complex to the activation of other signaling pathways such as STAT5 and JAK2. Noteworthy, t(6;9)-AMLs are characterized by activated STAT5 in presence of activated JAK2; Oancea et al., 2014), but here we showed that in t(6;9)-AML STAT5 activation is mainly regulated by SFKs and ABL1 and not by JAK2, similar to other cellular contexts (Ozawa et al., 2008; Silva, 2004).

The importance of a combination of network/cluster analysis, phosphoproteomics, and biochemical validation of each individual pathway to identify not only therapeutically active but also potentially undesirable consequences of drug effects is highlighted by the fact that all the above described signaling pathways except for JAK2 were important for the growth of DEK/CAN-positive cells. However in view of the inhibition of AKT/mTOR alone our data suggest that this may be clinically detrimental in the long term given the known role of STAT5 for LSC maintenance; Tam et al., 2013). This is of major importance not only in t(6;9)-AML, but also in breast cancer where similarly to our findings a direct interconnection between AKT/mTOR and STAT5 activation has been reported; Britschgi et al., 2012).

Although we provide evidence of a network of interacting CASPs in t(6;9)-AML (Figure 7H), the lack of functional interaction between SFK and ABL1, together with the need for both in the full activation of STAT5 does not allow the definition of one superordinate node. This seems to be unique for DEK/CAN-positive cells, because in most other tumors activated SFKs and ABL1 are functionally interconnected (Greuber et al., 2013; Khatri et al., 2016).

Our attempt to use highly selective allosteric ABL1 inhibitors to verify the impact of ABL1 activation was not successful, in contrast to findings in solid tumors, where activated ABL1 was effectively inhibited by allosteric inhibition; Wang et al., 2016). Possible reasons for these discrepancies of findings include differences in the activation process of ABL1, as indicated by the high rate of covalent post-translational modifications seen in DEK/CAN-positive cells.

The relevance of ABL1 in AML in general and in t(6;9)-AML in particular is still unclear. Activated ABL1 has a central role in a number of leukemias with activated kinases such as high risk Ph+ acute lymphatic leukemia (ALL) and a subset of Ph-like ALL which akin to t(6;9)-AML are leukemias with a high level of resistance. Although ABL1 activation shown in our work differs from that in these leukemias where it is translocation or gene fusion based it could have similar biological and clinical consequences (Boer et al., 2015; Roberts et al., 2012). Further studies should explore whether t(6;9)-AML may be a sort of Ph-like AML and may therefore be categorized together with the rare Ph+ AML.

In summary, we have devised an interactive proteomics approach as a powerful means of interrogating kinase signaling in leukaemia in a dynamic cellular context. Our novel SIP approach together with comparative phosphoproteomics move forward from the current strategy of mainly inhibiting a single activated pathway and suggests that simultaneous inhibition of selected kinase family members may be more effective in attenuating unwanted outputs in leukemia cells. Network analyses, as shown here, informs a specific selection of CASPs in each subtype of leukemia or even in each individual patient which should be targeted in order to maximize response and to increase the precision of any therapeutic intervention.

## Acknowledgements

The study was supported by a grant from German Cancer Aid (109787) to M.R. N.G. was supported by a fellowship from FAZIT-Foundation, Frankfurt, Germany.

## Author Contributions

Conceptualization, M.R. C.C.; Methodology, N.G., M.W., A.O., A.A.M., P.H., C.C.; Investigation, N.G., M.W., A.O., M.R., C.G., H.H., A.A.M., Formal Analysis, N.G. M.W., A.O., P.H., Ma.W., A.A.M., M.R., C.C.; Resources, N.G., M.W., M.R., C.C.; Writing - Original Draft, M.R., O.G.O., C.A., C.C.; Writing - Review & Editing O.G.O., C.A., M.R., C.C.; Funding Acquisition: M.R., O.G.O.; Supervision, D.B., K.J.H., M.R. C.C.

## Star Methods

### Contact for Reagent and Resource Sharing

Further information and requests for resources and reagents should be directed to and will be fulfilled by the Lead Contact, Claudia Chiriches.

### Experimental Model and Subject Details

#### Patient samples

Primary human AML samples were obtained with informed consent and used in agreement with the Declaration of Helsinki upon the approval of the local ethic committee of the Goethe University Frankfurt (approval number 329–10). Samples were maintained in X-vivo10 medium (Lonza) supplemented with 10% FCS (Hyclone/Perbio Science), with the addition of 20 ng/ml hIL-3, 50 ng/ml hSCF, 25 ng/ml hTPO and 50 ng/ml hFLT3-ligand (Miltenyi, Bergisch-Gladbach, Germany). For a CFU assay, freshly thawed bone marrow (BM) cells derived from a t(6;9)-positive patient were cultured in methyl-cellulose complete (MethoCult™ H4434, Stem-Cell Technologies) for 14 days and the colony number was determined in comparison to untreated samples.

### Mice and mouse models

All animal studies were performed in accordance with international animal protection guidelines and approved by the regulating institution for the animal facility respectively at the Goethe University Frankfurt (Regierungspräsidium Darmstadt - approval number F39/08) as well as at Cardiff University (HO – PIL number P1DBB264F). All mice were kept under standard conditions and diet and had a weight >20 grs.

#### Primary cell culture, colony forming unit assays (CFU), colony forming unit spleen-day 12 (CFU-S12)

For a CFU on murine leukemic cells freshly thawed BM cells from DEK/CAN-positive leukemic mice and healthy empty vector transduced control transplanted mice; Oancea et al., 2010) were plated in semi-solid medium supplemented with mIL-3 (20 ng/mL), mIL-6 (20 ng/mL) and mSCF (100 ng/mL)(Stem-Cell Technologies). On day 10 after plating, the number of colonies was determined. CFU-S12 assays were performed as described previously; Oancea et al., 2014). Briefly Sca1+/Lin-cells were isolated from the C57BL/6N mice as described and pre-stimulated for 2 days in DMEM medium with 10% FCS (Hyclone), 1% L-Glutamin, 1% Penicillin/Streptomycin, in the presence of mIL-3 (20 ng/mL), mIL-6 (20 ng/mL) and mSCF (100 ng/mL) (Cell Concepts) prior to retroviral transduction and cultivation for 10 days in DMEM medium supplemented with 10% FCS, mIL-6, mSCF, and mIL-3. The cells were then harvested and inoculated into lethally irradiated (11 Gy) female recipient mice. The inoculated mice were culled 12 days later, and the spleens were fixed in Tellysniczky’s fixative for colony counting.

Freshly thawed bone marrow cells derived from a t(6;9)-positive patient were cultured in semi-solid medium (MethoCult™ H4434, Stem-Cell Technologies) in the presence of the indicated compounds for 14 days and the colony number was determined in comparison to the untreated samples.

#### Syngeneic DEK/CAN-AML model

For induction of secondary AML 5×10^3^ thawed BM cells from DEK/CAN-positive leukemic mice were inoculated i.v. into sublethally (4.4 Gy) irradiated 6-8 weeks old female C57/BL6N mice (> 20g)(Charles River) via tail vein. Randomisation and experimental units were established. Starting from day 5 after injection 4 cohorts (n=7-8) were treated with solvent (PBS), dasa (20mg/Kg/day), BEZ (50mg/Kg/day) and dasa/BEZ in combination, respectively. After 21 days of treatment the mice were culled and spleen size determined.

### Cell lines and cell culture

All cell lines (293T, FKH-1, HEL, JURKAT, K562, Kasumi-1, RL, U937) were obtained from the German Collection of Microorganisms and Cell Cultures (DSMZ), Braunschweig, Germany. 293T cells were maintained in DMEM supplemented with 10% fetal calf serum (FCS) (Gibco, Thermo Fisher). The suspension cell lines were kept in RPMI-1640 medium supplemented with 10% FCS except for FKH-1 and Kasumi-1 cells which were maintained in 20% FCS. The following U937 clones were obtained from Martin Ruthardt’s laboratory: MTB45 (empty vector-control), PR9 (expressing PML/RARa), RUNX-1/ETO (expressing HA-RUNX-1/ETO), BCR/ABL; Puccetti et al., 2003) or DEK/CAN (expressing HA-DEK/CAN); Oancea et al., 2010). All these cells expressed their respective transgene under the control of MT-1 promoter inducible by 100 μM ZnSO4 (Zn^2+^); Ruthardt et al., 1997). These U937 clones were authenticated confirming the Zn^2+^-induced expression of the respective transgene by Western blotting (data not shown).

### I-Tasser

I-TASSER (Iterative Threading ASSEmbly Refinement)(https://zhanglab.comb.med.umich.edu/I-TASSER) was used to generate the hypothetic 3D structure of the coiled-coil domain in CAN based on its aa. sequence (Zhang, 2008). In a first step („threading“) the aa. sequence was compared to sequences present in the Protein Data Bank (PDB)(https://www.ebi.ac.uk/pdbe/); Burley et al., 2017) for which the 3D structure is known. In a second step („assembly“) the resulting structure fragments were assembled to create hypothetic 3D models. The energetically most favorable models were then analyzed in a last step („refinement“) for the presence of possible hydrogen-bonds and the most favorable backbone torsions simulations were calculated.

### Compounds

With the exception of PF-114 (kindly provided by Dr. Ghermes Chilov, Fusion Pharma, Moscow, Russia) all inhibitors used in this study were purchased from Selleckchem www.selleckchem.com). The 1000x stock solutions were obtained by dissolving the compounds in dimethyl sulfoxide (DMSO)(Sigma) then further diluted to working concentrations in the corresponding medium prior to use. For the colony assays the compounds were diluted directly in semi-solid medium.

### Plasmids, Oligos

The cDNAs encoding PML/RARα, RUNX-1/ETO or DEK/CAN and the related retroviral PINCO vectors have been previously described; Oancea et al., 2014). The DEK/CAN helix mutants were obtained using the Cold Fusion Cloning Kit (System Biosciences) using the following primers (5‘-3‘) DC Helix1-fw GCC AGT GCT AAC TTG GAT GTG AAT GAT GTT; DC Helix1-rev AAC ATC ATT CAC ATC CAA GTT AGC ACT GGC; DC Helix2-fw AAA TTT GCT GTC CAA CAA AGG CACG TGC TT; DC Helix2-rev AAG CAG GTG CCT TTG TTG GAC AGC AAA TTT; DC Helix3-fw CTG CTT GTG CCA GAG AGC CTG TCC TCG GCT; DC Helix3-rev AGC CGA GGA CAG GCT CTC TGG CAC AAG CAG; DC Helix4-fw AGT TTT GAC AGT GAC AAG ACC CCA CCA GTG; DC Helix4-rev CAC TGG TGG GGT CTT GTC ACT GTC AAA ACT; DC Helix5-fw GTG AGA TCC ACT GCT ACG TCC TGT AAA GAT; DC Helix5-rev ATC TTT ACA GGA CGT AGC AGT GGA TCT CAC. Point mutations in the putative GSK3 β phosphorylation sites were introduced by site directed mutagenesis using the QuikChange Site Directed Mutagenesis Kit (Agilent) according to the manufacturer’s instructions. The following primers were used: (5’-3’) DEK S30A-fw GTC CCA GAG AGG AGG CCG AGG AGG AAG AGG; DEK S30A-rev CCT CTT CCT CCT CGG CCT CCT CTC TGG GAC AAG CTG; DEK 240/241-rev CAG CTT CTT TAT CTT CAT CAT CTG CAG CCT CTT CCT TGT TTT TCT TTT C; DEK-A40G-fw AGG GAA CCC CCG CCC AGC CCG CG; DEK-A40G-rev CGC GGG CTG GGC GGG GTT CCC T; DEK-A196G-fw AAG AAA AAG TAG AGA GGT TGG CAA TGC AAG TCT CTT CCT TAC; DEK-A196G-rev GTA AGG AAG AGA CTT GCA TTG CCA ACC TCT CTA CTT TTT TCT T; DEK-T502G-fw GAG GTT CTT GAT TTG GAG AGA GCA GGT GTA AAT AGT GAA CTAG; DEK-T502G-rev CTA GTT CAC TAT TTA CAC CTG CTCT CTC CAA ATC AAG AAC CTC; DEK-T676G-fw ACC AAA TGT CCT GAA ATT CTG GCA GAT GAA TCT AGT AGT GAT G; DEK-T676G-rev CAT CAC TAC TAG ATT CAT CTG CCA GAA TTT CAG GAC ATT TGG T. The sequences were then transferred into the pEntry vector (Gateway – Thermo Scientific, Darmstadt Germany) for further recombinations into destinations vectors. The TAP-TAG DEK/CAN constructs were generated by transferring a TAP-tag sequence from a TAP-tag-pUC19 (kindly provided by Elena Puccetti, Institute for Molecular Biology and Tumor Research, Philipps University, Marburg, Germany) in frame with the sequences encoding the mutants in pEntry by using the Cold Fusion cloning kit (System Biosciences,). The correctness of sequence was confirmed for all constructs by Sanger sequencing.

### Interaction Proteomics

#### SILAC labeling and TAP-TAG purification

For a complete labeling cells were grown in SILAC medium (DMEM supplemented with 10% FCS dialyzed in order to eliminate the unlabeled a.a. (Gibco), 1% L-Glutamine, 1% Pen/Strep, essential amino acids) for at least 5 days or 5 divisions; Ong et al., 2002). Arginine and lysine (1%) enriched in stable isotopes of 13C and ^15^N (SILAC heavy) were used to label the cells before transfection with the empty vector control (TAP-), whereas the unmodified a.a. (SILAC light) were used for the cells to be transfected with a vector encoding TAP-DEK/CAN or TAP tagged mutants. Transfection of 293T cells was performed by calcium-phosphate precipitation according to widely established protocols. 48 hours later the cells were lysed and TAP-tagged proteins were precipitated in two consecutive steps, The eluted proteins from the heavy labeled control cells were mixed 1:1 with each of the eluted light-labeled sample proteins. The mixed protein samples were run on a SDS-gel, subjected to in-gel trypsin digestion and the resulting peptides. MS and MS/MS data were acquired with a LTQ-Orbitrap mass spectrometer (Thermo Fisher Scientific) online coupled to the LC system. The evidenced peptides were processed using the Proteome Discoverer™ Software 1.4 (Thermo Fisher Scientific) and raw data were searched against Uniprot Human Protein database, and a list based on proteins with at least 2 identified peptides was created. In addition only proteins with >70% of light labeled peptides were considered, subtracting the heavy labeled peptides as background.

#### Bioinformatic data elaboration: Interaction network analysis, Ingenuity^©^ Pathway Analysis (IPA)

Three different data bases for the analysis of potential functional interaction between proteins were used. STRING, functional protein association networks (http://string-db.org) allows to investigate functional protein-protein interactions, by representing the union of all possible protein–protein links; Jensen et al., 2009). The Biological General Repository for Interaction Datasets (BioGRID; https://thebiogrid.org) houses genetic and protein interactions curated from the primary biomedical literature for all major model organism species and humans which allows to investigate not only binary but also complex multi gene/protein interactions; Stark et al., 2006). IntAct (http://www.ebi.ac.uk/intact/) protein interaction database and analysis system for molecular interaction data; Kerrien et al., 2007).

### Phosphoproteomics

For phosphoproteomic analysis the FKH1 cells were splitted at a density of 250,000 cells /ml and allowed to rest overnight. The next day cells were treated for 6h with 20 nM Dasatinib, 1 μM Imatinib, 10 μM PP2 and 20 nM Torin1.

#### Tandem Mass Tag (TMT) Labeling and Phospho-peptide enrichment

Aliquots of 100μg of up to ten samples per experiment were digested with trypsin (2.5μg trypsin per 100μg protein; 37°C, overnight), labeled with TMT ten plex reagents according to the manufacturer’s protocol (Thermo Fisher Scientific) and the labeled samples pooled. This pooled sample was then desalted using a SepPak cartridge (Waters) and subjected to TiO2-based phosphopeptide enrichment according to the manufacturer’s instructions (Pierce). The phospho-enriched sample was evaporated to dryness and then resuspended in 1% formic acid prior to analysis by nano-LC MSMS using an Orbitrap Fusion Tribrid mass spectrometer (Thermo Fisher Scientific).

#### Nano-LC Mass Spectrometry

The TMT-labelled phospho-enriched sample was fractionated using an Ultimate 3000 nano-LC system in line with an Orbitrap Fusion Tribrid mass spectrometer (Thermo Fisher Scientific). In brief, peptides in 1% (vol/vol) formic acid were injected onto an Acclaim PepMap C18 nano-trap column (Thermo Scientific). After washing with 0.5% (vol/vol) acetonitrile 0.1% (vol/vol) formic acid peptides were resolved on a 250 mm × 75 μm Acclaim PepMap C18 reverse phase analytical column (Thermo Fisher Scientific) over a 150 min organic gradient, using 6 gradient segments (5-9% solvent B over 2min., 9-25% B over 94min., 25-60%B over 23min., 60-90%B over 5min., held at 90%B for 5min and then reduced to 1%B over 2min.) with a flow rate of 300 nl min^−1^. Solvent A was 0.1% formic acid and Solvent B was aqueous 80% acetonitrile in 0.1% formic acid. Peptides were ionized by nano-electrospray ionization at 2.0kV using a stainless-steel emitter with an internal diameter of 30 μm and a capillary temperature of 275°C.

All spectra were acquired using an Orbitrap Fusion Tribrid mass spectrometer controlled by Xcalibur 2.0 software (Thermo Fisher Scientific) and operated in data-dependent acquisition mode using an SPS-MS3 workflow. FTMS1 spectra were collected at a resolution of 120 000, with an automatic gain control (AGC) target of 200 000 and a max injection time of 50ms. The TopN most intense ions were selected for MS/MS. Precursors were filtered according to charge state (to include charge states 2-7) and with monoisotopic precursor selection. Previously interrogated precursors were excluded using a dynamic window (40s +/− 10ppm). The MS2 precursors were isolated with a quadrupole mass filter set to a width of 1.2m/z. ITMS2 spectra were collected with an AGC target of 5000, max injection time of 120ms and CID collision energy of 35%.

For FTMS3 analysis, the Orbitrap was operated at 60 000 resolution with an AGC target of 50 000 and a max injection time of 120ms. Precursors were fragmented by high energy collision dissociation (HCD) at a normalised collision energy of 55% to ensure maximal TMT reporter ion yield. Synchronous Precursor Selection (SPS) was enabled to include up to 5 MS2 fragment ions in the FTMS3 scan.

#### Data Analysis

The raw data files were processed and quantified using Proteome Discoverer software v2.1 (Thermo Scientific) and searched against the Uniprot Human database (134169 sequences) using the SEQUEST algorithm. Peptide precursor mass tolerance was set at 10ppm, and MS/MS tolerance was set at 0.6Da. Search criteria included oxidation of methionine (+15.9949) and phosphorylation of serine, threonine and tyrosine (+79.966) as variable modifications and carbamidomethylation of cysteine (+57.0214) and the addition of the TMT mass tag (+229.163) to peptide N-termini and lysine as fixed modifications. Searches were performed with full tryptic digestion and a maximum of 1 missed cleavage was allowed. The reverse database search option was enabled and all peptide data was filtered to satisfy false discovery rate (FDR) of 5%. The resulting phosphopeptides were used to generate the Heatmap in RStudio and for the comparative pathway analysis with IPA^©^ (Qiagen).

### Immunodetection

#### Immunoblotting

Immunoblot analyses were performed according to widely established protocols. The following kits and antibodies were used: mTor Substrates Antibody Sampler Kit, Phospho-Akt Pathway Antibody Sampler Kit, anti-pSTAT5-Y694, anti-pABL-Y245 and Y412, anti-pSRC-Y416, anti-SRC, and anti-pLCK-Y505, anti-pc-Myc-S62 (all from Cell Signaling). Anti-ABL, anti-STAT5, anti-c-Myc (Santa Cruz Biotechnology); anti-pLYN-Y396, anti-pc-Myc-T58; Y74 (Abcam), anti-LYN (BD Biosciences), anti-α-Tubulin (Lab Vision). Blocking and antibody incubation were performed in 5% low-fat dry milk (Carl Roth). Washing was performed in Tris-buffered saline containing 0.1% Tween20 (TBS-T) followed by incubations with secondary antibody coupled with horseradish-peroxidase for staining with enhanced chemoluminiscence substrate. Blots were “stripped” using “RestoreWestern blot Stripping Buffer” (Perbio Science). Imaging and elaboration was performed with the LI-COR Odyssey Fc system (LI-COR Biosciences).

#### Intracellular Flow Cytometry (ICF)

5×10^5^ cells/sample were fixed with Cytofix buffer (BD Biosciences) according to the manufacturer’s protocol. Permeabilization was performed with ice-cold 90% methanol for 30 min. Cells were incubated with the primary antibody pSTAT5-Y694-AlexaFluor 647 (BD Biosciences), pAKT-S473-PE-Vio770, pCRKL-Y207-APC (Miltenyi) and the non-labelled pABL-Y245 or IgG control for 40 min at RT. Washing was performed with PBS containing 1% FCS and 0.1% sodium azide. Fluorescence was measured immediately on a FACS Fortessa (BD Biosciences).

#### Confocal Laser Scan Microscopy (CLSM)

Cells were cultured over-night on poly-D lysine covered chamber slides (Falcon or Corning), washed with TBS (10 mmol/L Tris-HCl pH 8, 150 mmol/L NaCl), fixed in 4% paraformaldehyde (AppliChem) for 15 minutes, and permeabilized with 0.1% Triton-X in TBS. Blocking was performed in 3% (w/v) BSA (Sigma) and 0.1% Tween20 in TBS for 60 minutes. Cells were incubated with polyclonal rabbit anti–pABL (Y245)(Cell Signaling) or monoclonal anti–c-ABL (Santa Cruz). After extensive washing in 0.01% Tween20 in TBS, cells were stained with Alexa Fluor 647-conjugated goat anti-mouse or anti-rabbit Ig antibodies (Life Technologies). Nucleus staining was obtained using DAPI (Life Technologies). The slides were mounted with Moviol (Sigma). Images were acquired by a Leica TCS-SP5 confocal microscope (Leica Microsystems) under identical conditions for pinhole opening, laser power, photomultiplier tension, and layer number. Identical parameters were applied for all samples during data elaboration by Fiji software (www.fiji.sc). Final picture elaboration (cutting, contrast) was performed with GIMP-software.

### Proliferation/cytotoxicity

Proliferation was assessed by using the XTT proliferation kit according to the manufacturer’s instructions (Roche).

### Statistical analysis

Statistical significance (p<0.05) was determined using one-way ANOVA with Bonferroni posttest using GraphPad Prism 5.0 software (GraphPad Software).

## Supplementary Informations

**Supplementary Figure 1. DEK/CAN Mutants for the Subtractive Interaction Proteomics. A.** Expression of mutant construct in comparison to DEK/CAN. All constructs were fused in frame with a N-terminal HA-tag for the immunodetection by an anti HA-Ab. **B**. Influence of point mutations in the putative GSK3-sites alone and together with the mutated CKII sites on the overall phosphorylation of DEK/CAN. After Qiagen PhosphoProtein purification the different fractions were loaded on a SDS-PAGE. Nuclear localization was assessed by an anti-Histone 3 Ab (α-H3). The bars represent a quantification of the relative reduction of phoshorylation based on the intensity of the bands normalised to the H3 bands. **C.** Co-localization between DEK/CAN and its phosphorylation mutants by confocal laser scan microscopy. Phaco – phase contrast; green fluorochrome – GFP-DEK/CAN; red fluorochrome – Alexa 594-conjugated 2° Ab detecting the anti-HA Ab. **D.** Co-localization between DEK/CAN and its helix mutants. green fluorochrome – GFP-DEK/CAN; red fluorochrome – SNAP signal given by mutant construct fused to a N-terminal SNAP-tag.

**Supplementary Figure 2. Subtractive Proteomics – A. representative experiment.** Comparison between interaction partner of DEK/CAN and those of the mutants. The numbers expressed SILAC labeling degree of the DEK/CAN sample. All proteins in which a component was SILAC labelled (from empty vector transfected control cells) were taken as experimental noise and therefore excluded from further analysis. Orange: exclusive binder to DEK/CAN not found in any of the performed experiments for any of the mutants. Grey: although EBs in this experiment, these proteins were found to interact at least with one mutant in another experiment performed and therefore not considered as EBs. **B-H Graphical representation of STRING network analyses.** For all these analyses also the related IntAct URLs are given. **B.** RPS14 and RPS19. **C.** RAB1A and RAB6A. **D.** Clusterin (CLU). **E.** S100A7. **F.** PCBD1. **G.** Serpin B3. **H.** IDH3.

**Supplementary Figure 3 - Graphical representation of a BioGrid network analyses. A.** RPS19. **B.** RPS19. **C.** Clusterin (CLU). **D.** IDH3. **E.** Serpin B3. **F.** S100A7. **G.** RAB6A. **H.** RAB1A. **I.** RAB5A and RIN1 centered network with RAS and ABL1.

**Supplementary Figure 4 – Pathway analysis of p**hosphoproteomic data sets from FKH1 cells treated with Torin1, PP2, imatinib, and dasatinib – IPA (Ingenuity Pathway Analysis). Signaling pathways significantly (p> 0.05) influenced by the treatments and the relative numbers of peptides with increased (red bars) or decreased (green bars) phosphoralytion upon treatment as compared to untreated controls and consequent activation/inhibition of respective signaling pathways (orange – activation; blue – inhibition).

**Supplementary Figure 5 – SFK inhibition in FKH1 by pona(tinib). A.** Immunoblotting - The indicated antibodies detected phosphorylated and total SRC, LYN, and STAT5. Tubulin – loading control. U937, HEL and RL cells – controls. **B.** Proliferation - relative XTT values represent the mean of three independent experiments +/− SEM.

**Supplementary Figure 6 - JAK2 inhibition by ruxolitinib does not inhibit FKH1 cells**. HEL, Jurkat – positive and negative controls. Relative XTT values represent the mean of three independent experiments +/− SD.

**Supplementary Figure 7 - Induction of the signaling pathways is not caused by autocrine/paracrine activation through RTKs - A.** Inhibition profile of the indicated TKIs. **B.** The cytotoxicity/growth effect of the indicated TKIs on FKH1 cells was assessed by relative XTT values. Results for ruxolitinib, dasatinib, and ponatinib were taken from other figures in the paper. Here are presented mean values of one representative experiment out of three with similar results each performed in triplicates +/− SEM.

